# Asymmetric tethering by exocyst *in vitro* requires a Rab GTPase, a v-SNARE and a Sac1-sensitive phosphoinositide lipid

**DOI:** 10.1101/2023.08.08.552456

**Authors:** Guendalina Rossi, Gabrielle C. Puller, Patrick Brennwald

## Abstract

Tethering factors play a critical role in deciphering the correct combination of vesicle and target membrane for subsequent fusion. The exocyst plays a central role in tethering post-Golgi vesicles to the plasma membrane, although the mechanism by which this occurs is poorly understood. We recently established an assay for measuring exocyst-mediated vesicle tethering *in vitro* and we have adapted this assay to examine the ability of exocyst to tether vesicles in an asymmetric fashion. We demonstrate that exocyst differs from another post-Golgi vesicle tethering protein, Sro7, in that it is fully capable of tethering vesicles with functional Rab GTPase, Sec4, to vesicles lacking a functional Rab GTPase. Using this assay, we show that exocyst requires both the Rab and R-SNARE, Snc1, to be present on the same membrane surface. In contrast, using Sac1 phosphatase treatment, we demonstrate a likely role for phosphoinositides on the opposing Rab-deficient membrane. This suggests a specific model for exocyst orientation and its points of contact between membranes during heterotypic tethering of post-Golgi vesicles with the plasma membrane.

## Introduction

Tethering of transport vesicles to the appropriate membrane at the correct time and space is a critical component of generating and maintaining organelle and plasma membrane identity (Yu and Hughson, 2010). An important model for understanding the process of exocytic transport is the budding yeast, S. cerevisiae. In particular, it has been shown that vesicle tethering of post Golgi vesicles at the plasma membrane requires the Rab GTPase Sec4 and two direct effectors: the multisubunit exocyst complex and the tomosyn/Lgl homologs Sro7/Sro77 (Guo et al., 1999; Grosshans et al., 2006, Watson et al., 2015).

Exocyst is a member of the CATCHR family of multisubunit tethering complexes and consists of eight subunits: Sec3, Sec5, Sec6, Sec8, Sec10, Sec15, Exo70 and Exo84 (Terbush et al.,1996, Stanton and Hughson, 2023). The cryo-EM structure of the exocyst has shown that all the subunits have a similar structural organization and interact with one another to form an elongated structure 32nm long and 13nm wide (Mei et al., 2018). Initial contact between the exocyst and the vesicle surface is thought to involve the direct interaction between Sec4-GTP and the C-terminus of the Sec15 subunit of the exocyst which is present at one end of the elongated structure (Guo et al., 1999). The exocyst also interacts with the R-SNARE Snc1/2 through its Sec6 component (Shen et al., 2013) at a site which is located at the opposite end of the complex from Sec15. The dual requirement for both Sec4 and Snc places additional specificity onto post-Golgi vesicle tethering by the exocyst. An important component of exocyst tethering with the plasma membrane is the phosphoinositide PI(4,5)P2 which has been shown to bind to both the Sec3 and Exo70 subunits of the exocyst complex (He et al., 2007).

Sro7 is a member of the tomosyn/Lgl family of tumor suppressors. The crystal structure shows it is composed of two adjacent beta propellers which are conserved amongst all family members (Hattendorf et al., 2007). There is one binding site for Sec4 on the surface of Sro7 which maps to a conserved cleft between the two beta propellers (Watson et al., 2015). Homo-oligomerization of Sro7 has been shown to occur during vesicle tethering *in vitro* suggesting this is a mechanism by which Sro7 binds to Sec4 at the vesicle surface, bringing the separate membranes into proximity (Rossi et al., 2018). Exocyst and Sro7 both work in parallel and interact directly with one another though the Exo84 subunit of the exocyst (Zhang et al., 2008). Additionally, Sro7 and Rho GTPases, concentrated at sites of polarized growth, can activate exocyst tethering by increasing the strength of exocyst binding to Sec4 and Snc1/2 at the vesicle surface respectively (Miller et al., 2023). For both exocyst and Sro7, the initial docking event is thought to be followed by SNARE mediated fusion by promoting the localized assembly of SNARE monomers into fusion complexes at sites of polarized growth (Wu et al., 2008; Hattendorf et al., 2007).

We recently reconstituted an *in vitro* vesicle:vesicle tethering assay using purified Sro7 and exocyst protein and fluorescently-labeled post-Golgi vesicles isolated from yeast (Rossi et al., 2020). The *in vitro* assay closely mirrored tethering *in vivo* as both tethers relied on the presence of a functional Sec4 for vesicle tethering to occur. Additionally, the exocyst-mediated *in vitro* tethering also demonstrated a requirement for the R-SNARE Snc1/2 on the vesicle surface. When we modified the assay to include both Rab-proficient and Rab-deficient vesicles, we observed that Sro7-mediated symmetric vesicle tethering and functioned as a tether only when both opposing membranes contained Sec4 (Rossi et., 2018). This suggests Sro7 could function to help cluster, retain, and concentrate post-Golgi vesicles at sites of polarized growth in the cell. Here we show that unlike Sro7, the exocyst mediates asymmetric vesicle tethering between a Rab-proficient and a Rab-deficient surface. We have used this behavior to test several fundamental properties about how the exocyst recognizes each of these two membranes and how it is likely to be aligned during the tethering process. We demonstrate that the Rab and SNARE requirements for exocyst tethering are from the same membrane and not from opposing membranes while a Sac1-sensitive phosphoinositide, likely PI4P, is important solely on the opposing membrane. These results suggest exocyst is likely to adopt an alignment parallel to the surface of the vesicle during vesicle tethering in a manner that would be distinct from that suggested for the structurally related CATCHR family member Dsl1 (Stanton and Hughson, 2023).

## Results and Discussion

### Assaying the exocyst for asymmetric tethering activity

The membrane tethering process involves bringing two membranes into close proximity without fusion. In order to study post-Golgi vesicle tethering biochemically, we developed a reconstitution assay using isolated post-Golgi vesicles from yeast and purified forms of either of two vesicle tethering factors, Sro7 or the multisubunit exocyst complex (Rossi et al., 2015; Rossi et al., 2020). Both exocyst and Sro7 have been shown to be direct effectors of the Rab GTPase, Sec4 (Guo et al.,1999; Jin et al., 2011; Watson et al., 2015) and we have shown that functional Sec4 is required for both effectors to mediate vesicle tethering *in vitro* (Rossi et al., 2015; Rossi et al., 2020). While Sro7 is able to mediate *in vitro* vesicle tethering on its own, exocyst-mediated tethering shows a requirement for the presence of Sro7 in order to exhibit significant tethering activity *in vitro* (Rossi et al., 2020). We previously reported a simple modification of the *in vitro* assay which allowed us to determine if both membranes involved in a tethering reaction require a functional Sec4 protein on each of the two opposing surfaces or if the presence of Sec4 on one membrane is sufficient (Rossi et al., 2018). The assay involves using one population of vesicles which contains a functional GFP-Sec4 (green) derived from a *sec6-4* strain, while the other population of vesicles labeled with FM4-64 (red), lacks a functional Sec4, as it was derived from a *sec4-8* strain (Goud et al., 1988). As a positive control, we isolated FM4-64–labeled vesicles from a *sec6-4* strain, which results in (red) post-Golgi vesicles with wild-type Sec4 on their surface. The vesicles used in this assay were normalized by immunoblot analysis of vesicle marker proteins Snc1/2, Sso1/2 and Sec4 to make sure they were present in equal amounts in the tethering assay (Figure 1B). If Sec4 is required on only one of the two membranes in the tethering reaction, we would predict that the FM4-64– labeled vesicles obtained from the *sec4-8* mutant strain would be able to tether or “co-cluster” with the GFP-Sec4–labeled vesicles when mixed together in the presence of the tethering factor being examined. In contrast, if Sec4 is symmetrically required on each vesicle surface, for one vesicle to form a tether with another vesicle, then we would expect to see little or no detectable FM4-64 fluorescence in the GFP-Sec4– labeled clusters when *sec4-8* vesicles labeled with FM4-64 are used. However, we would predict that we would detect coincident fluorescence if FM4-64–labeled *sec6-4* vesicles (containing Sec4) are included instead of FM4-64 labeled *sec4-8* vesicles.

**Figure 1.**
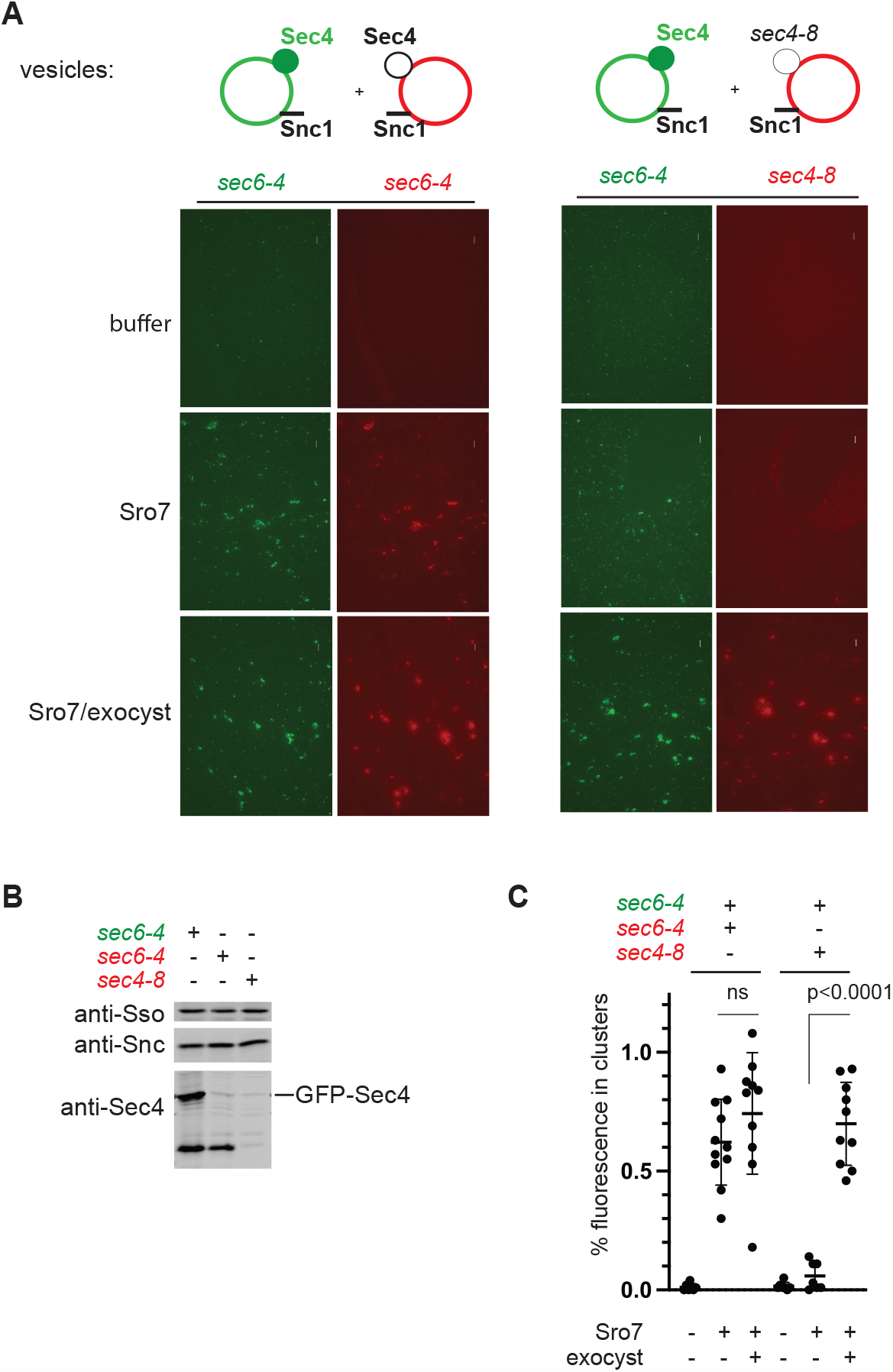
Asymmetric vesicle tethering by the exocyst. (A) Red vesicles labeled with the lipid dye FM4-64 were isolated from both *sec6-4* and *sec4-8* mutant strains and mixed with green vesicles isolated from a *sec6-4* mutant strain expressing GFP-Sec4 (*CEN*) in an *in vitro* tethering assay in the presence of Sro7 alone or Sro7 and wild type exocyst complex. Scale bar 5μm. (B)Immunoblot analysis of vesicle fractions used in the assay with vesicle marker proteins: Sso1/2, Snc1/2 and Sec4. (C) Quantitation of ‘mixed’ vesicle clusters defined as vesicle clusters in the TRITC channel which show 40% overlap with clusters in the FITC channel. Error bars represents SD obtained from counting images at 60x magnification. P values were obtained using a two-tailed Student’s t-test.

The results of these experiments are shown in Figure 1A, and quantitation is shown in Figure 1C. Vesicle tethering was quantitated using automated detection by commercially available software with a slight modification to the method we recently described (Miller et al., 2023; Materials and Methods). As we have seen previously (Rossi et al., 2018), the presence of Sro7 resulted in co-clusters only when Sec4 was present on both populations of vesicles (Figure 1A middle row). However, in contrast to Sro7-mediated tethering, when purified exocyst is included in the asymmetry assay we observe quite robust and significant co-clustering of FM4-64 labeled *sec4-8* vesicles (lacking functional Sec4) with GFP-Sec4 labeled vesicles (Figure 1A, C). This striking result may reflect an inherent property of the exocyst that is important for heterotypic post-Golgi vesicle tethering to the plasma membrane within the cell. This novel property of the exocyst in the *in vitro* tethering assay allowed us to test other fundamental aspects of how the exocyst might recognize and tether two opposing yet distinct membranes.

### Snc is required on the Sec4-proficient but not the Sec4-deficient vesicles in exocyst-mediated asymmetric tethering

We have previously shown that exocyst-mediated tethering requires both the Rab GTPase, Sec4, and the v-SNARE, Snc1/2 (Rossi et al., 2020). The asymmetric tethering assay described above allowed us to ask if the requirement for Snc1/2 is on the same membrane as that containing Sec4, on the membrane deficient for Sec4 (*sec4-8*), or both. To answer this question, we constructed modified versions of the yeast strains used above, where the only source of the R-SNARE, Snc1/2, was under a regulatable *GAL* promoter (*pGAL-SNC1*). Using these strains, we were able to generate both green Sec4-proficient vesicles which were depleted of Snc1 and red Sec4-deficient vesicles that were depleted of Snc1 (Figure 2B). The vesicles were normalized following immunoblot analysis and then mixed in a tethering assay with the respective red Sec4-deficient or green Sec4-proficient vesicles in the presence of both Sro7 and exocyst (Figure 2A,C). The results show clearly that Snc1 depletion of GFP-Sec4 vesicles causes a near total loss of asymmetric tethering with *sec4-8* vesicles (Figure 2C left graph). In contrast, Snc1 depletion of *sec4-8* vesicles results in robust tethering with GFP-Sec4 vesicles. In fact, quantitation of these results demonstrated a significant, nearly 3-fold, increase in asymmetric tethering of the Snc1-depleted *sec4-8* vesicles compared to the undepleted *sec4-8* control vesicles (Figure 2C right graph). These results demonstrate that during asymmetric tethering the recognition of R-SNARE and the Rab GTPase by exocyst occurs on the same membrane. The requirement for Sec4 and Snc1/2 on the same vesicle suggests a mechanism for how the tethering reaction might act as a proofreading mechanism to ensure that only vesicles with the appropriate combination of both Rabs and R-SNAREs are presented for tethering to the plasma membrane. The surprising increase in tethering when the R-SNARE was removed from the Rab-deficient membrane might reflect either the removal of a steric inhibition or an increase in properly oriented tethering complexes in the absence of any competing signals.

**Figure 2.**
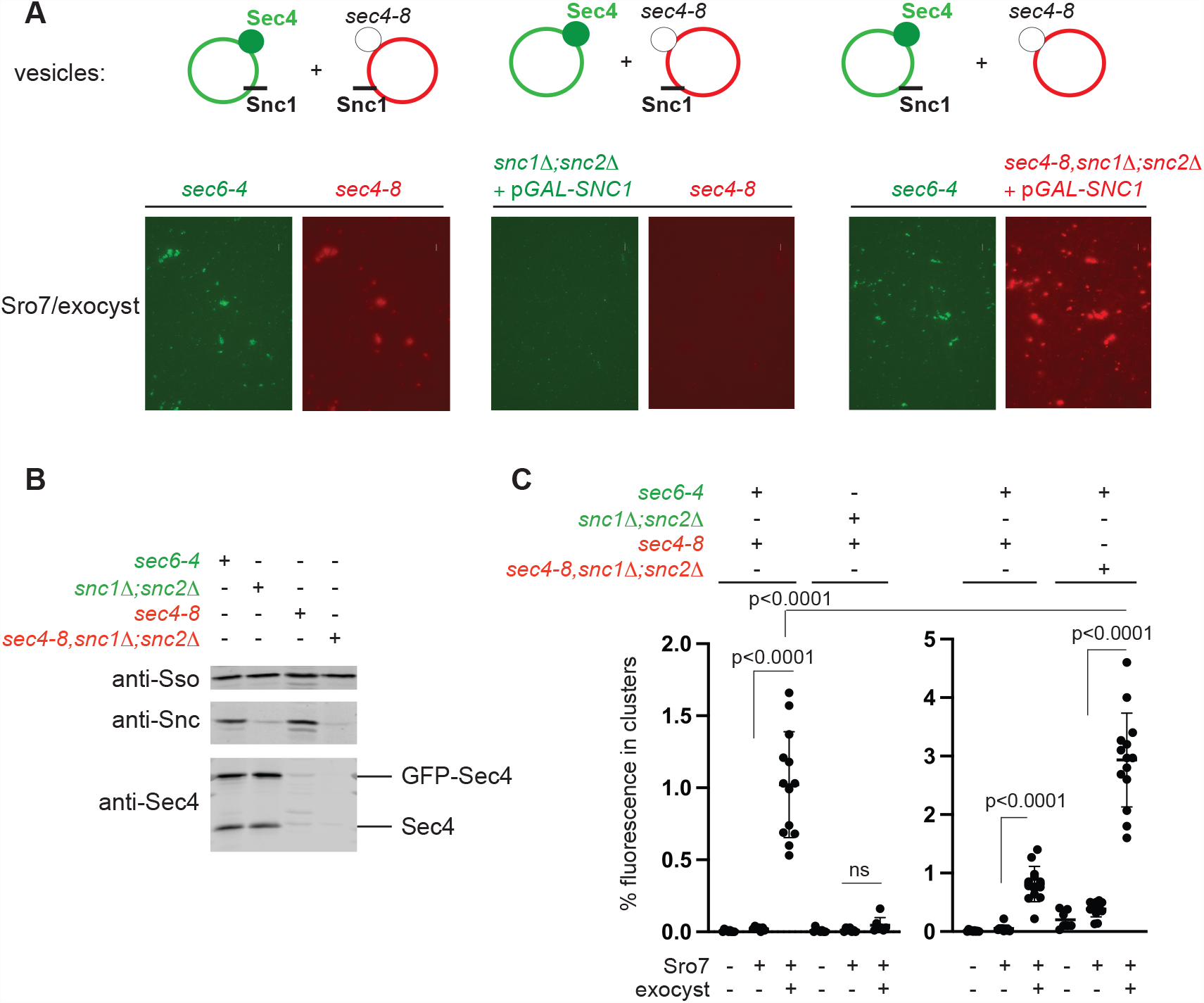
The R-SNARE Snc is required on the same membrane as Sec4 in asymmetric vesicle tethering by the exocyst. (A) Snc1/2 was depleted from green ‘Sec4-proficient’ vesicles expressing GFP-Sec4 or from red ‘Sec4-deficient’ vesicles containing the *sec4-8* mutation by genetic manipulation which involved placing Snc1 under a *GAL* promoter in strains expressing GFP-Sec4 (*snc1*Δ; *snc2*Δ + p*GAL*-*SNC1*) or strains containing a *sec4-8* mutation (*sec4-8, snc1*Δ; *snc2*Δ +p*GAL*-*SNC*1). Vesicles were then isolated, normalized and used in an *in vitro* tethering assay in the presence of Sro7 alone or Sro7 and wild type exocyst complex. Scale bar 5μm. (B) Immunoblot analysis of vesicle fractions used in the assay with vesicle marker proteins: Sso1/2, Snc1/2 and Sec4. (C) Quantitation of ‘mixed’ vesicle clusters defined as vesicle clusters in the TRITC channel which show 40% overlap with clusters in the FITC channel. Error bars represents SD obtained from counting images at 60x magnification. P values were obtained using a two-tailed Student’s t-test.

### A role for PI-4-phosphorylated lipids in asymmetric tethering by the exocyst

The results described above leave an open question of how exocyst is able to recognize and bind to the Sec4 and Snc-deficient vesicles during asymmetric tethering.

Phosphoinositide lipids are attractive candidates to perform this function. Phosphinositides have been implicated in exocyst function in post-Golgi transport; the Exo70 subunit has been shown to bind both PI(4,5)P2 and PI4P (He et al., 2007) while Sec3 has also been shown to interact with PI(4,5,)P2 (Zhang et al., 2008). Although PI(4,5)P2 is enriched at sites of polarized growth on the plasma membrane, only PI4P is thought to be present on the surface of post-Golgi vesicles (Santiago-Tirado et al., 2011; Mizuno-Yamasaki et al., 2010). To examine the role of phosphoinositides in our assay, we made use of the Sac1 enzyme, which has a well described phosphoinositide phosphatase activity. *In vivo* this enzyme is thought to act primarily on PI4P, but *in vitro* also shows activity on the monophosporylated phosphoinositides PI3P and PI5P (Guo et al., 1999, Zhong et al., 2012).

We treated fluorescently-labeled *sec4-8* and GFP-Sec4 vesicles with purified recombinant catalytically active Sac1^2-511^ or the catalytically inactive Sac1^2-460^ (Cai et al., 2014) as described in Materials and Methods. Following Sac1 treatment, vesicles were subjected to a final purification by pelleting though a sorbitol cushion and analyzed by immunoblot (Figure 3B and Figure 4B). We first analyzed the effect of Sac1 treatment on the red Sec4-deficient, *sec4-8*, vesicles in an assay mix with green (GFP) Sec4-proficient vesicles, Sro7 and exocyst. The results of this experiment, seen in Figure 3A and Figure 3C, demonstrate clearly that treatment of the red Sec4-deficient vesicles with the catalytically active Sac1^2-511^ dramatically inhibit the ability of the treated vesicles to tether to green untreated Sec4-proficient vesicles. In contrast, both the *sec4-8* vesicles treated with catalytically inactive Sac1^2-460^ and mock-treated vesicles demonstrate comparable tethering activity to each other (Figure 3A,C)-although both have a slightly lower activity than that seen in Figure 1.

**Figure 3.**
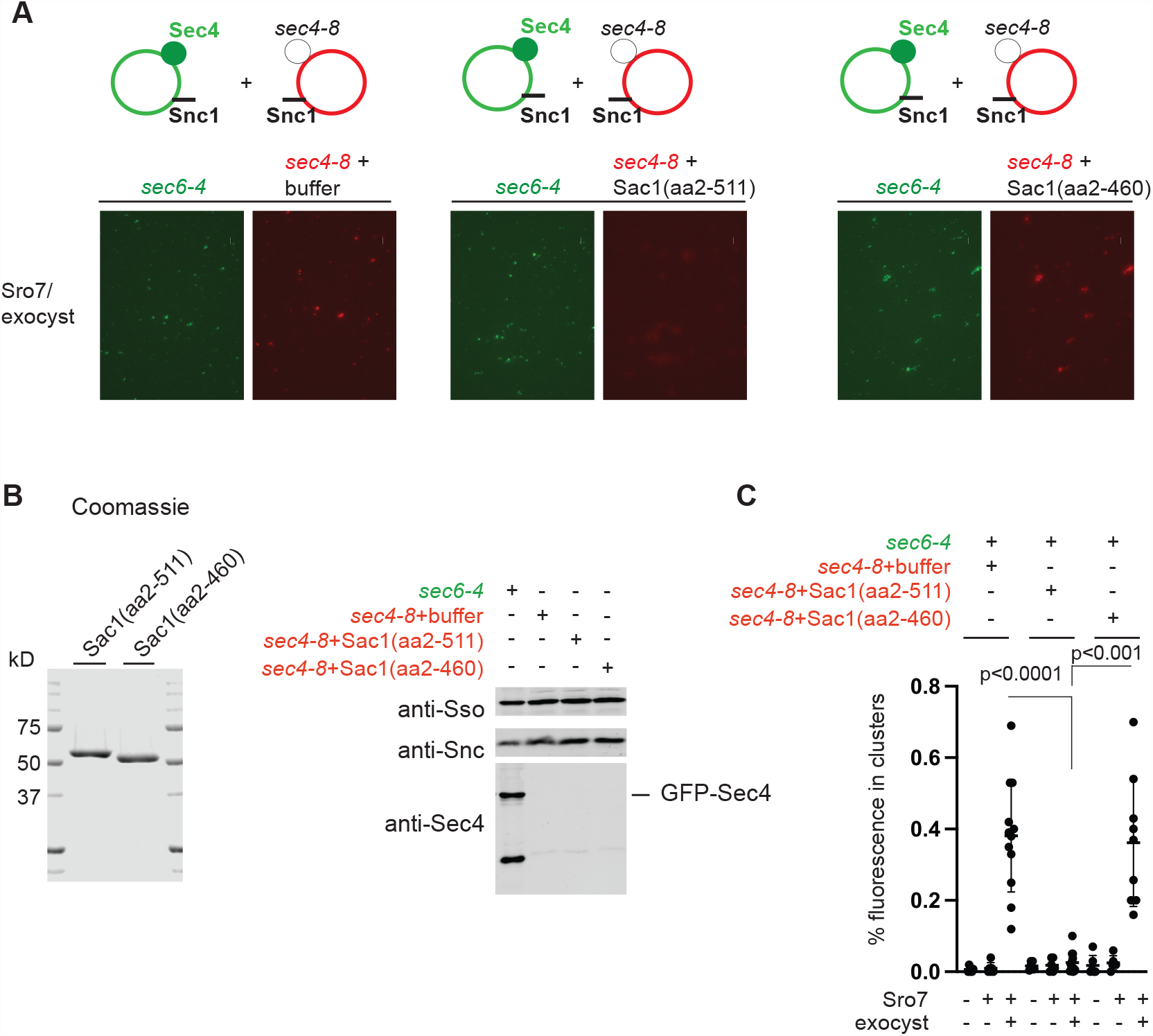
Sac1 treatment of the *sec4-8* vesicles disrupts asymmetric tethering by the exocyst. (A) Red ‘Sec4-deficient’ vesicles generated from a *sec4-8* mutant strain were treated with active Sac1(aa2-511), inactive Sac1(aa2-460) or buffer only before using in an asymmetric assay with green ‘Sec4-proficient’ vesicles, Sro7 and exocyst. (B) Immunoblot showing the normalization of the vesicles used in the assay with vesicle marker proteins: Sso1/2, Snc1/2 and Sec4. Coomassie of the purified Sac1 enzymes used in the assay is shown to the left. (C) Automated quantitation of ‘mixed’ vesicle clusters defined as vesicle clusters in the TRITC channel which show 40% overlap with clusters in the FITC channel. Error bars represents SD obtained from counting images at 60x magnification. P values were obtained using a two-tailed Student’s t-test.

**Figure 4.**
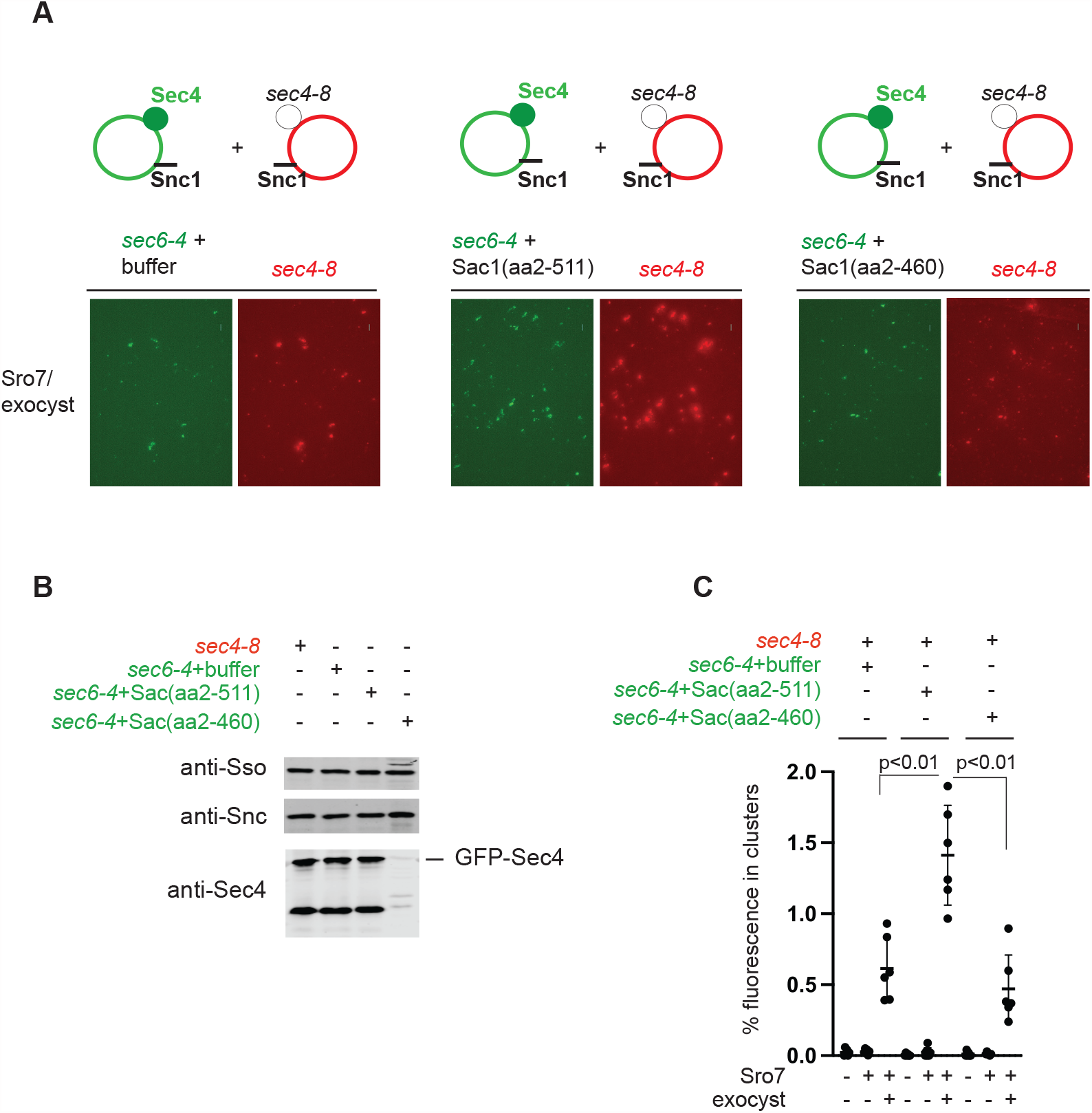
Sac1 treatment of GFP-Sec4 vesicles promotes asymmetric tethering by the exocyst. (A) Green ‘Sec4-proficient’ vesicles generated from a GFP-Sec4/*sec6-4* mutant strain were treated with active Sac1(aa2-511), inactive Sac1(aa2-460) or buffer only before using in an asymmetric assay with red ‘Sec4-deficient’ *sec4-8* vesicles, Sro7 and exocyst. (B) Immunoblot showing the normalization of the vesicles used in the assay with vesicle marker proteins: Sso1/2, Snc1/2 and Sec4. (C) Automated quantitation of ‘mixed’ vesicle clusters defined as vesicle clusters in the TRITC channel which show 40% overlap with clusters in the FITC channel. Error bars represents SD obtained from counting images at 60x magnification. P values were obtained using a two-tailed Student’s t-test.

To determine if phosphoinositides have a role in the activity of the Sec4-proficient vesicles used in the asymmetric tethering assay, we carried out a set of Sac1 treatments on the GFP-Sec4 vesicles identical to those used with the *sec4-8* vesicles above. The results of this experiment are shown in Figure 4A-C. In contrast to the results for the *sec4-8* vesicles, treatment of GFP-Sec4 vesicles with active Sac1^2-511^ caused an apparent increase--rather than a decrease--in tethering activity when compared to catalytically inactive Sac1^2-460^ or buffer controls (Figure 4A). In fact, quantitation of these results demonstrated a significant, roughly 2-fold, increase in asymmetric tethering of the Sac1 treated GFP-Sec4 vesicles compared to the two control-treated GFP-Sec4 vesicles (Figure 4C). This surprising increase in tethering when PI4P was removed from the Rab-proficient vesicles is remarkably similar to the effect of Snc depletion we saw in Figure 2. Likewise, this may also reflect an increase in properly oriented tethering complexes via a reduction of any competing signals from the GFP-Sec4 vesicles.

Taken together these experiments demonstrate a role for a phosphorylated phosphoinositide lipid species in exocyst-mediated tethering *in vitro*. PI4P, generated by Pik1 kinase in the Golgi, has been shown to be present on post-Golgi vesicles *in vivo* (Santiago-Tirado et al., 2011, Mizuno-Yamasaki et al., 2010) and is likely to represent the functional target of Sac1 treatment of post-Golgi vesicles in the assay. Furthermore, the specificity and asymmetry of the inhibitory and stimulatory effects of Sac1 treatment strongly support a role of phosphoinositide recognition in mediating the inherent asymmetry of the tethering of post Golgi vesicles with the plasma membrane. While PI4P appears to be the relevant lipid in the asymmetry assay *in vitro*, PI(4,5)P2 is the likely target of exocyst recognition of the plasma membrane in heterotypic tethering of vesicles *in vivo* (see below).

While PI4P is the primary target of Sac1 phosphatase *in vivo, in vitro* Sac1 has been shown to also act on PI3P and PI5P species (Zhong et al., 2012). PI5P is thought to be a short-lived species and not commonly detected in yeast (Foti et al., 2001) and PI3P is found on endosomal membranes (Hasegawa et al., 2017). Therefore, it is highly likely that the effects we see of Sac1 enzymatic activity in the tethering assay are mediated by loss of PI4P (conversion of PI4P to PI) from the vesicle surface.

### Models for exocyst-mediated membrane tethering

The results described in the previous sections have important mechanistic implications for models of how exocyst acts to tether two distinct membranes. Our results also suggest a model for the relationship between asymmetric tethering in this assay *in vitro* and heterotypic tethering of post-Golgi vesicles to the plasma membrane *in vivo*. First, the finding that the v-SNARE, Snc1/2, is required on the same membrane as the Rab GTPase, Sec4, (see Figure 2) has very important implications for structure-based models on how the exocyst complex is likely to be arranged during tethering. This is due to the fact that the binding sites for Snc1/2 and Sec4 have been shown to reside on distinct ends of the exocyst complex (Mei et al, 2019; Figure 5A). The site of Snc1/2 binding is known to reside within the Sec6 subunit or the “Sec6 cap” present at one end of the complex (Shen et al., 2013), while the binding site for Sec4 GTPase, is known to reside in the Sec15 subunit or “Sec15 pole” at the opposite end of the complex (Lepore et al., 2018, Wu et al., 2005). Simple modeling of the two possible arrangements of the exocyst cryo-EM structure with the Snc-depletion data demonstrate that the “extended tethering” model is inconsistent with our analyses. Rather, the “parallel tethering” model is in strong agreement with all of our experimental results. This immediately poses a new question. How does the exocyst recognize and physically interact with the Sec4-deficient, *sec4-8* vesicles during asymmetric tethering? An attractive answer comes from our studies with the Sac1 phosphoinositide phosphatase.

**Figure 5.**
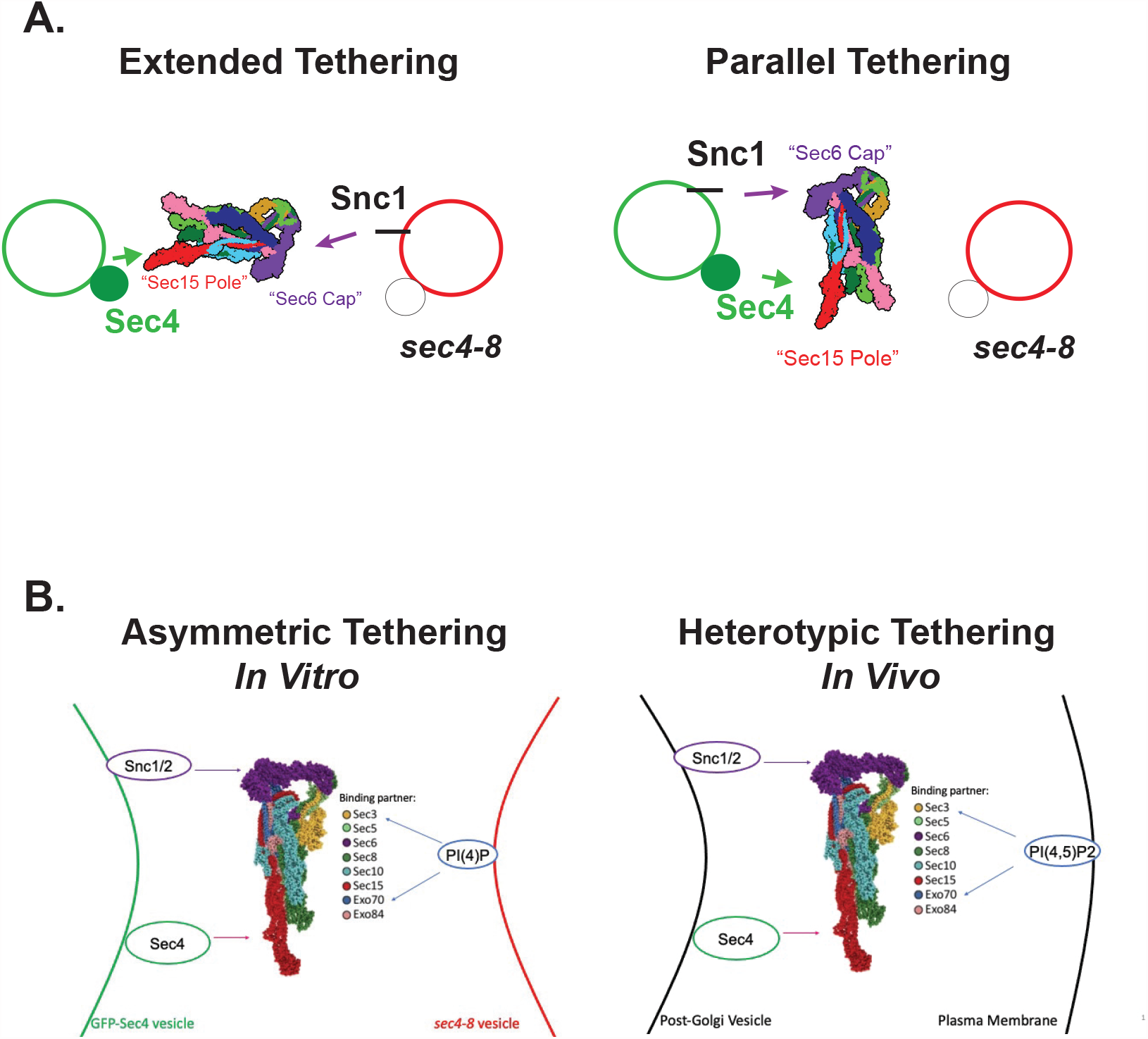
Models for exocyst-mediated tethering to the plasma membrane. (A) The cryo-EM structure of the exocyst places the binding sites for Sec4 and Snc1/2 on the opposite ends of the elongated exocyst complex. This allows for two distinct arrangements of the complex during tethering. The left shows what we refer to as “extended tethering” model where the Rab GTPase (Sec4) binding and R-SNARE (Snc1) binding occur on two distinct vesicle surfaces during tethering. The right shows “parallel tethering” where Rab and R-SNARE binding to the exocyst are provided from the same vesicle surface. (B) Comparison of asymmetric tethering of post-Golgi vesicles in the *in vitro* assay to heterotypic tethering of post-Golgi vesicles to the plasma membrane in the cell.

The demonstration that Sac1 treatment of post-Golgi vesicles has a potent inhibitory effect in exocyst-mediated asymmetric tethering suggests a role for phosphoinositides in this process. This fits nicely with the identification of PI(4,5)P and PI4P binding to the Exo70 subunit of the exocyst and the importance of this binding to the function of the exocyst *in vivo* (He et al., 2007). The Sac1 phosphatase is well known to have a high degree of specificity for phosphoinositide lipids *in vivo* and *in vitro* (Foti et al., 2001; Guo et al., 1999). Therefore we suggest that PI4P is likely to be the relevant phosphoinositide present on *sec4-8* vesicles in the *in vitro* system, as this lipid has been shown to be present at least transiently on post-Golgi vesicles, while PI(4,5)P2 presence is thought to be specific to the plasma membrane (Mizuno-Yamasaki et al., 2010; Ling et al., 2012). Importantly, we find that the inhibitory effect of Sac1 phosphatase treatment is quite specific to the Sec4-deficient, *sec4-8* membranes and has no effect on Sec4-proficient membranes. This suggests a simple model, seen in Figure 5B, where PI4P in the asymmetric tethering system *in vitro* plays a role analogous to the role of PI(4,5)P2 in heterotypic tethering of post-Golgi vesicles with the plasma membrane *in vivo*. Moreover, it suggests a simple mechanism where recognition of plasma membrane-specific phosphoinositides would play an important role in providing both the specificity and physical interaction with the “target” membrane during heterotypic vesicle tethering.

Maib and Murray (2022) have recently shown PI(4,5)P2 dependence in exocyst-recruitment to vesicle membranes *in vitro* using reconstituted mammalian exocyst and synthetic lipid bilayers. Interestingly in this system PI(4,5)P2 appears to be important on both the vesicle and target membrane whereas in our assay phosphoinositides do not appear to play a critical role on the Rab/vSNARE-containing vesicles. Future studies will be directed at determining whether this represents a difference between the assay systems--the synthetic lipids used by Maib and Murray lacked bo Rab or vSNARE components for example--or a difference between how yeast and mammalian forms of the exocyst utilize phosphoinositides in heterotypic vesicle tethering to the plasma membrane.

Recently, Hughson and colleagues have described an overall structural similarity between the exocyst and another CATCHR family MTC, the Dsl1 complex that mediates tethering of COP1 vesicles to the endoplasmic reticulum (Stanton and Hughson, 2023). This complex is thought to capture vesicles in an extended arrangement with one end binding vesicles and the other end associated with Q-SNAREs on the endoplasmic reticulum. Although the structural similarity of Dsl1 complex with exocyst is striking, the proposed arrangement for tethering appears to be quite distinct from what we have found in the present work. One difference in the systems is that the *in vitro* assay described here lacks an active t-SNARE component as the Sec9 Q-SNARE is completely absent from the reaction. Therefore, it is possible that other arrangements of the complexes may come into play during the trans-SNARE assembly step that may bring this structural similarity into play as the two membranes come into closer proximity.

## Materials and Methods

### Protein purification

Sro7 and exocyst purification were carried out as described in Rossi et al., 2015 and Miller et al., 2023.

Active Sac1(BB2586) was cloned as a BamHI-NdeI fragment (aa 2-511) into the pET15b vector. Inactive Sac1(BB2603) was subcloned as a NdeI-BamHI fragment (aa2-460) in the pET15b vector.

### Generation of vesicles labeled with FM4-64 or GFP-Sec4 for the *in vitro* asymmetry tethering assay

Post-Golgi vesicles labeled with FM4-64 were isolated from single *sec6-4, sec4-8* mutant strains and a double *sec4-8, snc1*Δ; *snc2*Δ+p*GAL*-*SNC1* secretory mutant strain. The *sec6-4* mutant strain was grown in YP+2% glucose overnight at the permissive temperature of 25^°^C to a final OD_599_of 0.6. Cells were then shifted to 37^°^C for 2 h to accumulate vesicles. Sodium azide was added to a final concentration of 20mM and 300 absorbance units were centrifuged and washed with ice-cold 10mM Tris, pH 7.5 and 20mM sodium azide. Cells were spheroplasted in 10ml of buffer (0.1M Tris, pH 7.5, 1.2M sorbitol,10mM sodium azide, 21 mM β-mercaptoethanol, and 0.05mg/ml Zymolyse 100T) for 30 minutes at 37^°^C and then lysed in 4 ml of ice-cold buffer (10mM triethanolamine, pH 7.2, and 0.8M sorbitol) containing protease inhibitors (2μg/ml leupeptin, 2μg/ml aprotinin,2μg/ml antipain, 14μg/ml pepstatin A and 1mM phenylmethylsulphonyl fluoride). The lysate was spun at 4^°^C to pre-clear unbroken cells and then spun at 30000xg_max_ for 15 min in a Sorvall centrifuge to preclear large membranes. Approximately 3 ml of lysate was then labeled on ice for 10 min with FM4-64 at a final concentration of 1μg/ml. The labeled lysate was the layered over a 2 ml ice cold sorbitol cushion (20% wt /vol in 10mM triethanolamine, pH 7.2) and spun at 128000g_max_ for 1 h at 4^°^C. The final high-speed pellet was resuspended in 600μl of lysis buffer and frozen at -80^°^C. For vesicles isolated from the *sec4-8* mutant strain the following modifications were made: 600 absorbance units were spheroplasted in 15 ml of buffer before lysis in 4 ml of lysis buffer and final resuspension in 1ml of lysis buffer.

Vesicles isolated from the *sec4-8, snc1*Δ; *snc*2Δ deletion strain containing a (*CEN*) plasmid expressing Snc1 from a *GAL* promoter were generated by growing cells in YP+3% raffinose and 1% galactose for 5 h to a final OD_599_ of 1.5 before shifting to YP+2% glucose for 15 h at 25^°^C to inhibit Snc1 expression from the *GAL* promoter. Cells were then shifted to the restrictive temperature of 36^°^C for 2h. 350 absorbance units were then spheroplasted in 10 ml of buffer, lysed in 4 ml of lysis buffer and resuspended in final volume of 400μl before freezing at -80^°^C.

Post-Golgi vesicles labeled with GFP-Sec4 were generated from both a *sec6-4* mutant strain containing a *CEN* plasmid expressing GFP-Sec4 and from a *snc1*Δ; *snc2*Δ secretory mutant strain expressing p*GAL*-*SNC1* (*CEN*) and GFP-Sec4(*CEN*). Vesicles isolated from the *sec6-4* mutant strain expressing GFP-Sec4 were obtained by growing the strain overnight at 25^°^C in synthetic media to an OD_599_ of 0.6 and then in YP+2% glucose for 1 h at 25^°^C before shifting the cells for 2 h to 36^°^C. 350 absorbance units were then spheroplasted with 10ml of buffer before lysis with 4ml of lysis buffer and final resuspension in 1ml of lysis buffer. Vesicles isolated from a *snc1*Δ, *snc2*Δ deletion strain expressing Snc1 under a regulatable *GAL* promoter (p*GAL*-*SNC1 CEN*) and expressing GFP-Sec4 from a *CEN* plasmid were grown in synthetic media with 3% raffinose and 1% galactose for 5 h to a final OD_599_ of 1.5 before shifting to YP+2% glucose for 17 h at 25^°^C to repress the *GAL* promoter, inhibit Snc1 expression and accumulate vesicles. 350 absorbance units were then spheroplasted in 10ml of buffer before lysis in 4ml of lysis buffer and final resuspension of vesicles in 1ml of lysis buffer. Vesicles were then frozen at -80^°^C.

Sac1 treated *sec4-8* or *sec6-4* vesicles were obtained by generating the vesicles as described above and then adding 50nM Sac1 (aa2-511), Sac1(aa2-460) or buffer only (10mM Tris 7.5, 150mM sodium chloride) with the vesicles for 1hr at 25^°^C. The vesicles were then diluted in 2.5ml of lysis buffer before layering on a 20% (v/v) sorbitol cushion in 10mM triethanolamine, pH 7.2. The samples were then spun at 128000g_max_ for 1 h at 4^°^C before resuspending in 450 and 350μl of lysis buffer respectively. Vesicle fractions were frozen at -80^°^C.

### Strain construction

BY3215 (*mat a, sec4-8::NatR, snc1Δ:URA3, snc2Δ:ADE8, pGAL-SNC1*) was generated by transforming BY101 (*mat a, snc1Δ:URA3, snc2Δ:ADE8, pGAL-SNC1)* with an integrating form of *sec4-8* linked with a *NatR* gene. Nourseothricin (Nat) resistant transformants were confirmed to be temperature-sensitive and demonstrated a requirement for galactose in growth media. Immunoblot analysis demonstrated loss of both Snc and Sec4 following shift to glucose media at 36 ^°^C. BY3212 (*mat a, snc1Δ:URA3, snc2Δ:ADE8, pGAL-SNC1, GFP-SEC4-pRS315*) was created by transforming BY101 with a *CEN/LEU2* plasmid expressing *GFP-SEC4*.

### *In vitro* asymmetry tethering assay

Vesicles generated as described above were thawed, spun in a cold microfuge for 2 min and then treated with MgCl_2_(3mM) and GTP*γ*S(1mM) for 60 min on ice. Vesicles were then mixed in equal amounts in the presence of Sro7(0.4μM) or Sro7(0.4μM) and exocyst (20nM) for 60 min at 30^°^C. Vesicle-vesicle clustering was detected by taking six images at 60x magnification in both FITC and TRITC channels and quantitated by automated detection with Imaris Software v9.9. Clusters in the TRITC channel that were above 1μm in diameter and which showed over 40% overlap with clusters in the FITC channel were quantitated as described in Miller et al., 2023 except that co-localization was obtained by using the object-object function in Imaris and the FITC fluorescence was set at a threshold of 10.

## Acknowledgements

We thank Wendy Salmon and the UNC Hooker Imaging Core Facility for help with the use of Imaris software in the automated quantitation of tethering in this study, and Benjamin Twara for technical help with the recombinant Sac1 purifications. This work was supported by National Institutes of Health award R01 GM054712 (to P. B). G.C.P. was supported in part by a grant from NIGMS award T32 GM135128. Wendy Salmon and the UNC Hooker Imaging Core Facility are supported in part by P30 CA016086 Cancer Center Core Support Grant to the UNC Lineberger Comprehensive Cancer Center.

